# A Closer-to-Brain Heterosynaptic Learning Rule for Spatiotemporal Spike Pattern Detection with Low-Resolution Synapse

**DOI:** 10.64898/2026.04.19.719429

**Authors:** Shunta Furuichi, Takashi Kohno

## Abstract

The brain is believed to process information efficiently in a different manner from deep learning-based artificial intelligence (AI). Brain-like next-generation AI is gaining attention owing to its potential to perform human-like, highly adaptive, robust, and power-efficient computation. To realize such AI, one crucial approach is the bottom-up implementation of the neuronal systems, capturing their electrophysiological characteristics in electronic circuits. However, this neuromorphic approach generally focuses on simplified neuronal models that do not refer to many biological findings. Developing closer-to-brain models is a natural direction that serve as a fundamental computing model for next-generation AI. One of the constraints of neuromorphic circuits is the bit resolution of synaptic efficacy memory, as the memory footprint scales with it precision. Although low-resolution synaptic efficacy is essential for minimizing memory circuit footprint and energy consumption, it generally leads to performance degradation in many tasks such as the spatio-temporal spike pattern detection. This study proposed a closer-to-brain learning rule that incorporates heterosynaptic plasticity (HP) induced by glutamate spillover. It is demonstrated that our model mitigates the performance degradation associated with low-bit resolution synaptic efficacy, achieving the pattern detection success rate with 3-bit resolution synaptic efficacy, which is comparable to 64-bit floating-point precision. Furthermore, the findings of the study indicate that HP based model accelerates the convergence of the synaptic effcacy and effectively potentiates the synapses relevant to the pattern detection while suppressing irrelevant ones, thereby promoting a bimodal distribution of synaptic efficacies. These findings may provide a basic framework for constructing an energy-efficient, brain-like next-generation AI that maintains high performance under hardware constraints.

## 1. Introduction

In recent years, artificial intelligence (AI) technologies based on deep learning have advanced rapidly, supported by improvements in the power of computing systems and the availability of big data. However, these technologies still have several limitations, including enormous power consumption during training and difficulties in continual and one-shot learning. Furthermore, many current deep learning-based AI systems are optimized for specific tasks. Adapting them to different tasks requires retraining or even redesigning the network architecture [1]. In contrast, the brain can execute a vast number of different tasks adaptively with a few experiences. To overcome the aforementioned limitations, efforts to build a brain-like next-generation AI are gaining importance [2].

An approach to develop AI systems inheriting the advanced computing ability of the brain is to build an artificial copy of the brain at the cellular and synaptic level. Although the field of neuromorphic engineering originally aligned with this direction, it currently focuses on implementing deep learning-based AIs with higher power efficiency. The importance of this bottom-up approach remains a crucial option for next-generation brain-comparable AI [3]. Owing to the intricate network of neuronal cells in the brain and the numerous aspects still to be clarified, developing a complete replica of the brain’s neural networks is unfeasible. Thus, the basic strategy is to describe the neuronal cells and synapses using tractable models. Generally, minimally simplified spiking neuronal network (SNN) models, comprising networks of integrate-and-fire-based (I&F-based) point neuron models connected by single- or double-exponential synapse models, have been utilized. For the plasticity rule, pair-based spike timing-dependent plasticity (STDP) rules have been adopted. To realize brain-comparable systems, it is crucial for expanding such baseline models closer to the brain by referencing biological data such as the electrophysiological properties of neurons and synapses, synaptic plasticity, and the connectome. Implementation of SNN models on dedicated electronic circuits [4, 5, 6] is also crucial from the application and scientific viewpoints. It is expected to contribute by “analysis by construction” to elucidate the information processing properties of the neuronal cells’ network. In these silicon neuronal networks, the current main technology for storing synaptic efficacy is the digital memory circuit, whose footprint and power consumption exponentially increase with its bit resolution.

This work proposes a novel STDP model that considers heterosynaptic plasticity induced by glutamate spillover (i.e. leakage of glutamate outside the synaptic cleft) [7, 8], which facilitates the use of the lower-bit-resolution memory for synaptic efficacy. The proposed model is incorporated into a single-neuron model with a leaky I&F spiking mechanism and a double-exponential synapse model [9]. The original work [9] suggested that an asymmetric STDP rule leads the neuron to fire in response to a specific spatiotemporal input spike pattern hidden within the Poisson spike trains. This model was selected as the baseline model for evaluation of our model because it is one of the simplest models composed only of biologically supported submodels, and the input spike trains consider the synaptic noise without assuming rate coding. These points are crucial because the proposed model is intended as a “closer-to-the-brain” model, which may provide some implication to the effect of glutamate spillover in the brain.

From an engineering point of view, reducing the bit-resolution of the synaptic efficacy memory is important because the circuit area and the power consumption of the digital memory storing synaptic efficacy increases with bit resolution. However, the use of low-resolution synaptic efficacy typically leads to a degradation in learning accuracy. In the previous efforts [10, 6], a learning rule called adaptive STDP has been proposed. The adaptive STDP is a model in which the time window that induces long-term depression (LTD) in asymmetric STDP is adjusted during learning. The adaptive STDP can mitigate the degradation of pattern detection performance in the baseline model [9]. However, a remaining problem is that the adjustment mechanism of the LTD window is independent of neural activity, whose biological support is not provided, rendering it less biologically plausible.

This work aims to overcome this limitation by referring to biological findings. As explained previously, the model is induced by glutamate spillover, and it can suppress the task performance degradation with lower bit resolution of synaptic efficacy. Here, the task is the spatiotemporal spike pattern detection in the baseline model [9], rather than the Modified National Institute of Standards and Technology (MNIST) classification, which is generally used in more “AI”-oriented works [11]. In this task, input spike trains in this task assume neither rate coding nor specific data properties (for example, image data). The task is more appropriate for closer-to-brain models because temporal coding is believed to be used in cortical neurons [12], and a single cortical neuron does not classify image data directly. We demonstrated that our model efficiently promotes the potentiation of synapses crucial for pattern detection and reduces the proportion of intermediate synaptic efficacy, thereby mitigating the degradation in pattern detection accuracy with 3-bit resolution synaptic efficacy. The remainder of this paper is organized as follows. In Section 2, the details of the methods used in this study are described. Section 3 presents the results, demonstrating the performance of our model and analyzing it. Finally, Section 4 concludes the paper.

## 2. Methods

### 2.1. Neuron and synapse model

The model of the neuronal dynamics follows that of a preceding work [9]. It comprises a leaky integrate-and-fire (LIF) neuron and a double-exponential synapse models expressed by spike response model (SRM). In this model, membrane potential *v* at each time step is represented by the sum of the excitatory postsynaptic potential (EPSP) kernel *ε* and a spike kernel *η*, as follows.

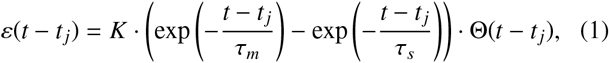

where K is a scaling constant, which is set so that the maximum amplitude of *ε* driven by a single spike equals to 1. *t* _*j*_ is the occurrence time of a presynaptic spike. *τ*_*m*_ and *τ*_*s*_ are the membrane and synaptic decay constants, respectively. Θ is the Heaviside step function, defined as 1 for *t* ≥ *t* _*j*_ and 0 otherwise. The spike kernel *η* describes the dynamics of the postsynaptic spike generation.

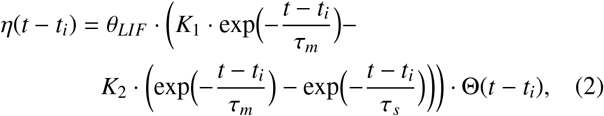

where *θ*_*LIF*_ is the threshold for spike gneration and *t*_*i*_ is the time of postsynaptic spike. *K*_1_ and *K*_2_ are the scaling constants tuning the amplitude of the membrane potential at firing a spike. Membrane potential *v* at each time step is described by the following equation:

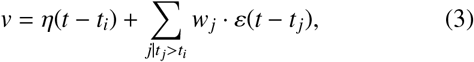

where *w*_*j*_ is the synaptic efficacy at synapse *j*. The parameter values of the neuron model are listed in Tables 2 and 3.

### 2.2. Spike timing dependent plasticity (STDP)

The standard learning model in this manuscript follows the preceding work [9]. The magnitude and direction (potentiation or depression) of the synaptic efficacy change are exponentially dependent on the time difference between pre- and post-synaptic spikes, as shown in Figure 1a and expressed in following equations:

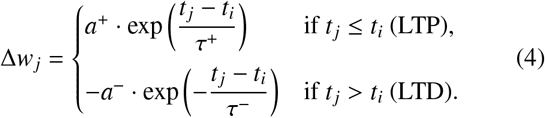

**Figure 1:**
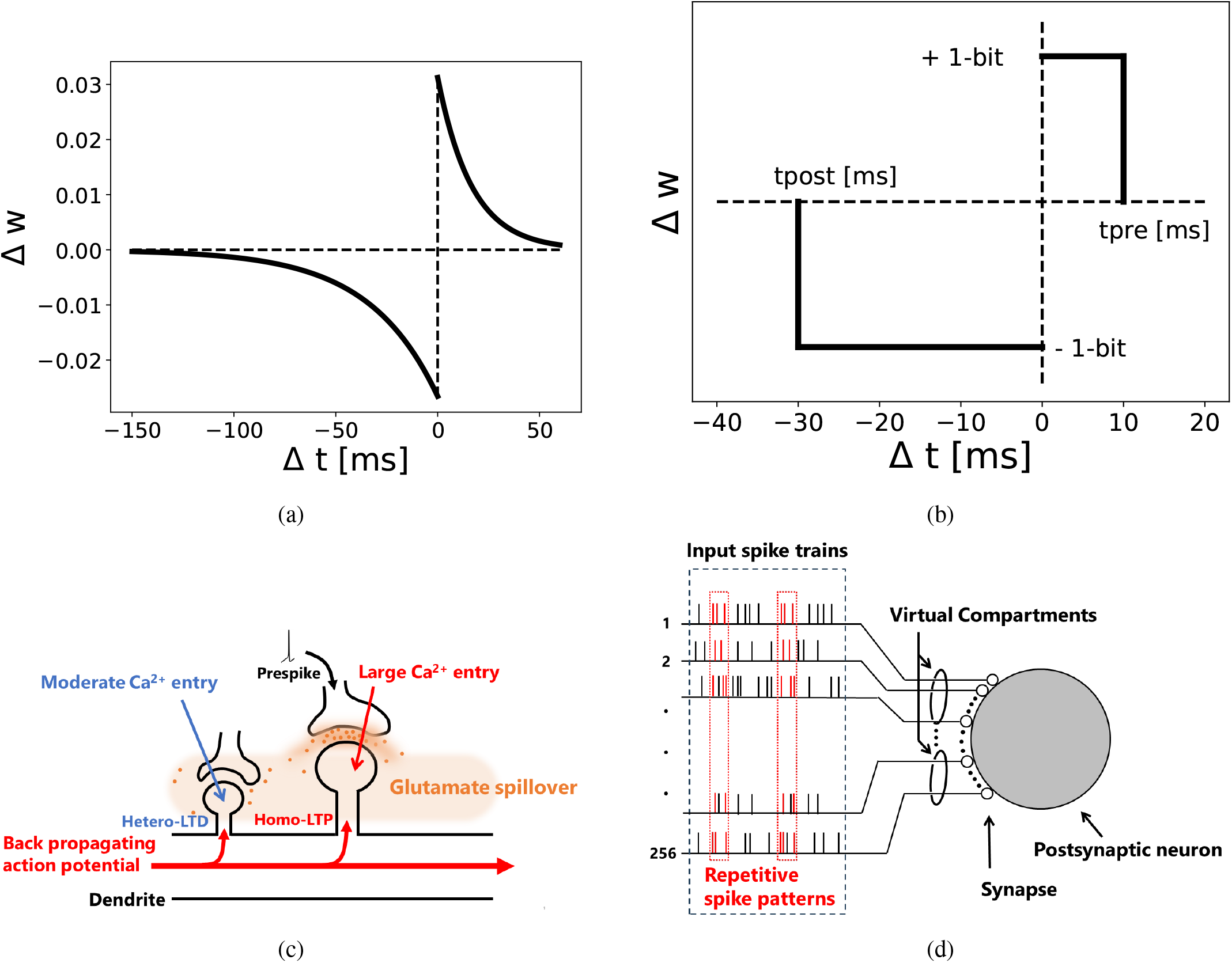
**a** Learning window of STDP. **b** A learning window of 1-bit STDP. **c** Mechanisms of heterosynaptic plasticity induced by glutamate spillover. When a presynaptic spike reach presynaptic terminal, glutamate (orange) is released into synaptic cleft and diffuse from extrasynaptic space. Deporalization at postsynaptic spine occurs by backpropagating action potential with certain concentration of extrasynaptic glutamate leads to moderate Ca^2+^ entry and heterosynaptic LTD (blue). **d** Overview of spike pattern detection model. A postsynaptic neuron recieves synaptic input via 256 synapses. Repetitive spike patterns (red) were embedded in noise. Virtual compartment was introduced to consider the diffusion of glutamate for heterosynaptic plasticity.

Here, Δ*w*_*j*_ is the amount of the synaptic efficacy change at each time step. *a*^+^ and *a*^−^ are the maximum values of the synaptic efficacy change. *τ*_+_ and *τ*_−_ are the time constants for the LTP and LTD windows in the STDP curve. In this work, we set synaptic efficacy change for LTP or LTD to zero when difference of pre- and post-spike times surpass 7 · *τ*^+^ and 7 · *τ*^−^, respectively.

For the learning model with discretized synaptic efficacy, we used the rectangular STDP for the learning rule [10, 6]. It is a simple version of the STDP, as shown in Figure 1b and expressed by following equations.

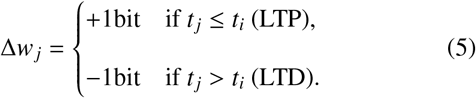

In the rectangular STDP learning rule, the synaptic efficacy change is 1-bit for each synaptic event. On the one hand, LTP induction results in a 1-bit increase in synaptic weight. On the other hand, LTD induction results in a 1-bit decrease in synaptic weight. The parameter values of the STDP are listed in Tables 2 and 3.

### 2.3. Glutamate spillover mediated heterosynaptic LTD

Glutamate is the principal excitatory neurotransmitter in the mammalian brain. Although glutamate was traditionally considered to be confined to the synaptic cleft for point-to-point transmission, recent evidences indicate that glutamate can diffuse into the extrasynaptic space. This phenomenon, known as glutamate spillover, can enable the activation of N-methyl-D-aspartate receptors (NMDARs) on neighboring cells [13, 8]. Based on these findings, we developed a model in which the extrasynaptic glutamate resulting from spillover activates NMDARs. In this study, we modeled the heterosynaptic LTD based on the following mechanism: when the concentration of the extrasynaptic glutamate reaches a certain threshold and coincides with postsynaptic depolarization via a backpropagating action potentials (bAP), the Mg^2+^ block is removed from NMDARs. This enables glutamate binding to trigger a Ca^2+^ influx. Because moderate Ca^2+^ influx preferentially induces LTD [14, 15], this process results in heterosynaptic LTD.

To integrate our heterosynaptic plasticity model into aforementioned neuronal model, the relative location between synapses on the dendrite has to be defined. Here, we introduced a “virtual compartment” in which the synapses are virtually grouped and considered as a member of a spatially closed synapse cluster. The glutamate concentration resulting from spillover is assumed to be homogeneous in each virtual compartment. In this study, we fixed the number of synapses in each virtual compartment throughout the simulation. The number of synapses for virtual compartment *i*, denoted as *S*_*i*_, was independently sampled from a uniform distribution *U*(*S*_*min*_, *S*_*max*_), where *S*_*min*_ and *S*_*max*_ represent the minimum and maximum number of synapses per compartment, respectively. We selected 6 and 10 for *S*_*min*_ and *S*_*max*_, respectively. The dynamics of the glutamate concentration in each compartment are governed by the following equation:

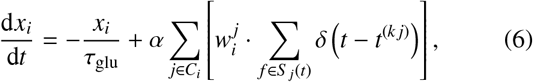

where *x*_*i*_ denotes the abstract concentration of extrasynaptic glutamate in virtual compartment *i, α* is the scaling constant, and *k* is the spike index at synapse *j*. Function *δ* is the Dirac delta function, defined such that it is zero for *t* − *t*^(*kj*)^ ≠ 0 and its integral over the real line is unity. *C*_*i*_ is the set of indices for synapses in virtual compartment *i*, and *S*_*j*_(*t*) is the set of spike indices until time *t* at synapse *j. τ*_*glu*_ is the decay time constant of extrasynaptic glutamate and calculated according to the following equations:

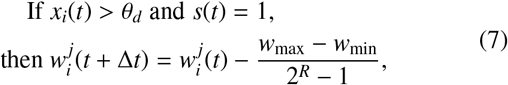

where *θ*_*d*_ represent the threshold of glutamate concentration for induction of heterosynaptic LTD. Function *s*(*t*) is 1 if a postsynaptic spike occurs, otherwise 0. *w*_*max*_ and *w*_*min*_ are maximum and minimum values of synaptic weight, respectively. *R* is the resolution of synaptic efficacy, set to 3 in this work. According to [8], the induction of LTP promotes glutamate spillover by driving withdrawal of perisynaptic astroglia. We incorporated this effect by considering the modification of decay time constant of extrasynaptic glutamate. This modification is represented by the following equation:

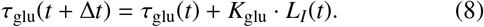

Here, *K*_*glu*_ is the modification rate of the decay time constant of extrasynaptic glutamate. *L*_*I*_(*t*) represents the number of LTP induction events in virtual compartment *i*. In this study, the model incorporated only the increasing dynamics of *τ*_glu_, as its decay time constant was beyond the time window considerd in this study [8]. We represent initial value of *τ*_glu_ as 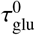. The parameter values of this model are listed in Tables 2 and 3.

### 2.4. Spatiotemporal spike pattern detection task

The model was evaluated using a spatio-temporal spike pattern detection task defined in [9]. This study demonstrated that a LIF neuron receiving inputs from synapses equipped with STDP can learn to detect repetitive spike patterns embedded in noisy spike trains. An overview of the spatio-temporal spike pattern detection model used in this study is shown in Figure 1c. The model consists of a single postsynaptic neuron and 256 excitatory synapses connecting to it. The input spike trains were generated following the methodology in [9]. First, background spike trains were generated as independently inhomogeneous Poisson process with an instantaneous firing rate randomly fluctuating between 0 Hz and 90 Hz. The maximum rate of the firing rate change was restricted to change from 0 to 90 Hz in 50 ms. Subsequently, a 50 ms bin within the background spike trains was randomly selected and the spikes in this bin were used as the spatio-temporal spike pattern to be detected (it is referred to as the target pattern hereafter). Subsequently, 10% of the 50 ms bins in the background spike trains were randomly chosen and the spikes in these bins were replaced by the target pattern. When we considered a noise, aditional 10 Hz noisy spikes and jitter noise was added to all spike trains throughout the simulation period. The jitter noise applied to the repetitive spike pattern was modeled as a Gaussian random variable with zero mean and standard deviation of 1 ms. The learning objective of this model is for the postsynaptic neuron, through STDP, to selectively fire only during the presentation periods of the target pattern. To quantify the performance, we first evaluated whether the spike pattern detection task was successful by two evaluation metrics: “Hit rate” and “Number of false alarms”. The hit rate was defined as the fraction of bins with the target pattern in which at least one postsynaptic spike occurred. The false alarm was defined as the postsynaptic spikes fired outside the bins without the target pattern. Both metrics were calculated for the time period after the evaluation start time till the simulation end time. Following the previous studies [9, 10, 6], we defined the successful pattern detection as the simultaneous satisfaction of two criteria: (1) the hit rate is 96% or higher, and (2) the number of false alarms is 0. Subsequently, the success rate was calculated by dividing the number of successful detections by the total number of pattern detection tasks performed using input spike trains generated with different random seeds.

In this work, four simulation setups were adopted (see Table 1). The number of synapses was set to 256 for all setups. On the one hand, a target pattern was coded in all 256 synapses in Setups 1 and 3. On the other hand, in Setups 2 and 4, the repetitive spike pattern was embedded in only half of the synapses. The input spike trains for Setups 3 and 4 were generated by applying the 10Hz and jitter noises. We performed simulations on the following three types of models with the four setups:

- 64-bit model: This model comprises 64-bit floating-point synaptic efficacy (the same as the original model [9] with different parameter values).
- 3-bit model: This model comprises synaptic efficacy discretized to 3-bit resolution, with the learning rule remained solely 1-bit STDP, similar to that in [10, 6].
- 3-bit-HP model: It is a 3-bit model expanded by the proposed heterosynaptic plasticity (HP).

**Table 1:** Simulation setups.

| Setup | $N_{syn}^{active} / N_{syn}$ | Noise |
| --- | --- | --- |
| 1 | 256 / 256 | NO |
| 2 | 128 / 256 | NO |
| 3 | 256 / 256 | YES |
| 4 | 128 / 256 | YES |

For all models and setups, the total simulation time was 150 s, and the evaluation start time was 120 s. In all simulation runs, the LIF model was computed using the SRM, and the glutamate dynamics within the virtual compartment were calculated using the fourth-order Runge-Kutta method. The simulation time step Δ*t* was set to 10 *µ*s.

### 2.5. Bayesian optimization (BO)

The performance of spike pattern detection is highly dependent on the model parameters. To improve the fairness of the model performance comparison, we introduced BO, a type of machine-learning-based black-box optimization method for tuning the parameters. In BO, we aim to maximize an objective function *f* : 𝒳 → ℝ, where 𝒳 is the parameter search space, and *r* ∈ ℝ is the success rate. In this study, the set of parameters that defines 𝒳 depends on the model, as follows:

- 64-bit model: (*τ*_*m*_, *θ*_*LIF*_, *τ*_+_, *τ*_−_);
- 3-bit model: (*τ*_*m*_, *θ*_*LIF*_, *t*_*pre*_, *t*_*post*_);
- 3-bit-HP model: (*τ*_*m*_, *θ*_*LIF*_, *t*_*pre*_, *t*_*post*_, 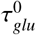, *K*_*glu*_, *θ*_*d*_).

Objective function *f* is the success rate for 50 trials with different input spike trains. The search space is listed in Table 3. The range of each parameter was as follows:

- *τ*_*m*_ : According to the experimental studies [16, 17, 18, 19], the value of the menbrane time constant is on the order of tens of milliseconds. Futhermore, a previous study [20] suggested that the optimal *τ*_*m*_ for pattern detection is on a time scale significantly shorter than the pattern duration. As the time length of the target pattern used in this study was 50 ms, the search range for *τ*_*m*_ was set to 2–15 ms.
- *θ*_LIF_ : We observed that the neuron generated no spikes when *θ*_LIF_ was 70 or higher because total EPSP could not reach the threshold. Thus, 70 was selected for the upper bound. To avoid highly frequent firing, the lower bound of the parameter was set to 20.
- *τ*_+_ : Experimental studies have shown that the width of the time window for inducing LTP is on the order of tens of milliseconds [21]. Based on this evidence, we set the search range to 10–30 ms.
- *τ*_−_ : The time window for LTD tends to be wider than that for LTP [21]. Therefore, we set the parameter search range to 10–100 ms.
- *t*_*pre*_ : We empirically found that *t*_*pre*_ larger than 10 ms tends to incur excessive firing. Therefore, we set the search range to 2–10 ms.
- *t*_*post*_ : We set the search range to 2–12 ms because the time window for LTD tends to be wider than that for LTP [21].
- 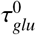 : The range of 10–100 ms was selected, because an experimental study reported that the lifetime of extrasynaptic glutamate is on the order of tens of milliseconds [13].
- *K*_*glu*_ : The search range was set to 0.0–0.1 ms. The upper bound (0.1 ms) ensures that *K*_*glu*_ is smaller than the initial value of 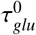, consistent with the minor withdrawal of astrocytes expected from a single LTP event. The lower bound (0.0) accounts for the possibility that this change occurs on a considerably longer timescale than other neural processes.
- *θ*_*d*_ : Because the value of *θ*_*d*_ remains experimentally undetermined, we defined the search range based on the model constraints. Given an initial synaptic weight of 0.286 as used in this study, a single presynaptic spike generates an extrasynaptic glutamate level of 0.000286 because *α* equals to 0.001. We reasoned that a single spike should be insufficient to trigger h-LTD; therefore, we set the threshold at 0.001, roughly one order of magnitude higher than the single spike contribution.

**Table 2:**
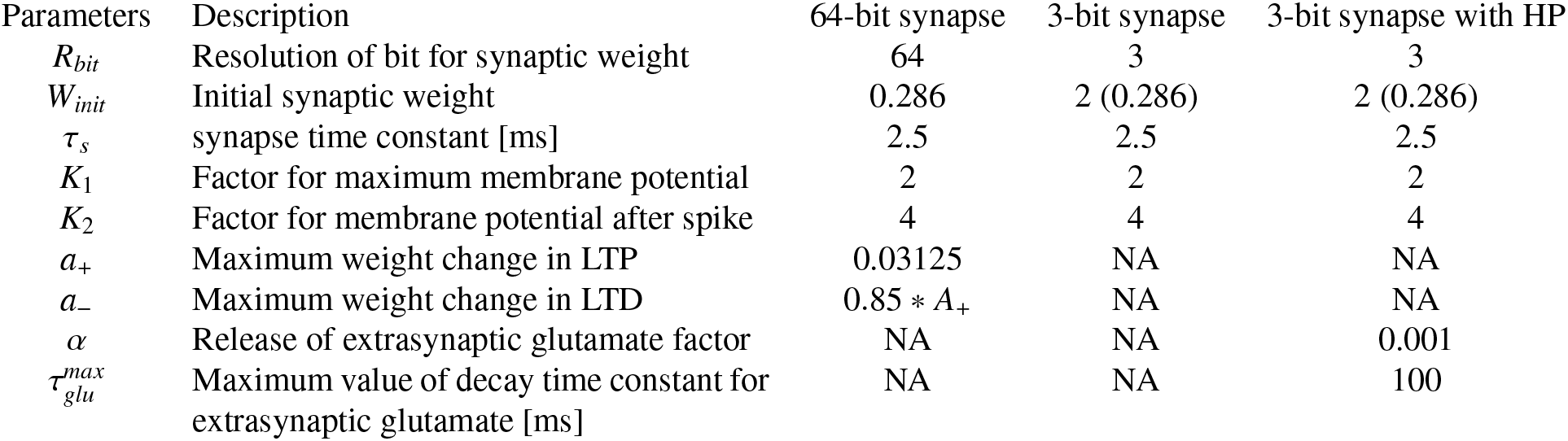
Fixed simulation parameters for neuron, synapse, synaptic plasticity.

**Table 3:** Parameter search ranges of Bayesian Optimization for neuron, synapse, and synaptic plasticity.

| Parameters | Description | 64-bit synapse | 3-bit synapse | 3-bit synapse with HP |
| --- | --- | --- | --- | --- |
| $\tau_m$ | Membrane decay time constant [ms] | (2, 15) | (2, 15) | (2, 15) |
| $\theta_{LIF}$ | Threshold of LIF model | (20, 70) | (20, 70) | (20, 70) |
| $\tau_+$ | Time constant for LTP window [ms] | (10, 30) | NA | NA |
| $\tau_-$ | Time constant for LTD window [ms] | (10, 100) | NA | NA |
| $T_{pre}$ | Spike time difference for LTP [ms] | NA | (2, 10) | (2, 10) |
| $T_{post}$ | Spike time difference for LTD [ms] | NA | (2, 12) | (2, 12) |
| $\tau_{glu}^0$ | Baseline of decay time constant for ex-trasynaptic glutamate [ms] | NA | NA | (10, 100) |
| $\tau_{glu}^{rate}$ | Change rate of decay time constant for extrasynaptic glutamate [ms] | NA | NA | (0.0, 0.1) |
| $\theta_d$ | Threshold of glutamate to induce heterosynaptic LTD | NA | NA | (0.001, 0.01) |

In each optimization step, a set of parameters was first selected from the parameter ranges defined in Table 3. The objective function, the success rate, was then calculated using these parameter values. It was computed as the average value over 50 simulation runs, each generated with a different random seed. The entire optimization process comprised 200 steps. We used hyperparameter tuning framework Optuna [22].

## 3. Results

### 3.1. Heterosynaptic LTD mitigate the degradation of pattern detection performance induced by low-resolution synaptic efficacy

The pattern detection success rates for each model and the corresponding model’s parameters obtained by the BO process (see Sec. 2.5) are listed in Tables 4, 5, 6, and 7. Across all models, the success rates were lower in Setup 3 compared to Setup 1, and in Setup 4 compared to Setup 2. One reason for this performance degradation is the jitter noise added to the target patterns, which disrupts the precise timing of postsynaptic spikes relative to the pattern presentation. The second reason is the addition of 10 Hz background noise to the input spike train. This increases the probability of false positives, where coincidental noise inputs trigger firing events even in the absence of the repetitive spike pattern. Additionally, comparing Setup 1 with Setup 2, and Setup 3 with Setup 4, we observed a degradation in success rates across all models. It is the same tendency as in [10, 6]. This can be attributed to the reduced number of synapses coding the repetitive spike pattern. Reducing synaptic efficacy resolution from 64-bit to 3-bit resulted in significant performance drops of 34%, 36%, 30%, and 56% for setups 1 through 4, respectively. This substantial degradation is likely caused by the increased granularity of synaptic efficacy updates associated with lower bit resolution. Specifically, the coarse step size implies that a single plasticity event induces a large discrete change in synaptic efficacy. If LTD occurs at a synapse relevant to pattern detection or LTP occurs at a synapse irrelevant to pattern detection, it becomes difficult for the postsynaptic neuron to continue firing inside the target pattern and discontinue firing outside it, thereby severely disrupting the learning process. However, by incorporating HP into the 3-bit model, pattern detection success rates improved by 38%, 28%, 36%, and 10% for setups 1 through 4, respectively. This improvement is attributed to HP that can rapidly modify the synapses where an undesirable synaptic efficacy update was induced.

**Table 4:**
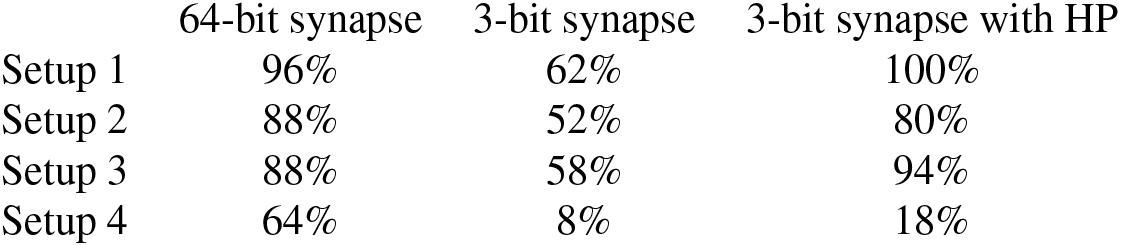
Results of the success rate in Bayesian optimization (BO) across different models and setups. Each value represents the best success rate of pattern detection achieved via BO. The objective function of the BO was the success rate, which was calculated from 50 trials with different input spike trains at each iteration.

|  | 64-bit synapse | 3-bit synapse | 3-bit synapse with HP |
| --- | --- | --- | --- |
| Setup 1 | 96% | 62% | 100% |
| Setup 2 | 88% | 52% | 80% |
| Setup 3 | 88% | 58% | 94% |
| Setup 4 | 64% | 8% | 18% |

**Table 5:** Parameters decided by Bayesian optimization in 64-bit synapse model.

| Parameters | Setup1 | Setup2 | Setup3 | Setup4 |
| --- | --- | --- | --- | --- |
| $N_{syn}^{active}/N_{syn}$ | 256/256 | 128/256 | 256/256 | 128/256 |
| Noise | NO | NO | YES | YES |
| $\tau_m$ | 0.0031948863467893152 | 0.003440148973420971 | 0.002 | 0.007302597740853504 |
| $\theta_{LIF}$ | 26.44303762019582 | 29.32144975424349 | 24.351756374386746 | 50.15201887114384 |
| $\tau_+$ | 0.02279718917051777 | 0.019339758328754664 | 0.020360043731346405 | 0.022148596142315572 |
| $\tau_-$ | 0.1 | 0.042686016044869525 | 0.0768496295009337 | 0.05325373839164086 |

**Table 6:** Parameters decided by Bayesian optimization in 3-bit synapse model.

| Parameters | Setup1 | Setup2 | Setup3 | Setup4 |
| --- | --- | --- | --- | --- |
| $\tau_m$ | 0.0036986293065501283 | 0.002 | 0.004478350300690541 | 0.0022805143283112354 |
| $\theta_{LIF}$ | 30.021831491539572 | 20.0 | 39.88905264515554 | 20.65591903337566 |
| $T_{pre}$ | 0.0026711630114835704 | 0.003261958745568283 | 0.0029916465901218946 | 0.002 |
| $T_{post}$ | 0.012 | 0.0111663884974305 | 0.009358482411180676 | 0.007490797596669883 |

**Table 7:** Parameters decided by Bayesian optimization in 3-bit synapse model with HP.

| Parameters | Setup1 | Setup2 | Setup3 | Setup4 |
| --- | --- | --- | --- | --- |
| $\tau_m$ | 0.0038845132198768524 | 0.0021945944304043938 | 0.004241670105649584 | 0.0037989868111908435 |
| $\theta_{LIF}$ | 34.445815909613614 | 20.0 | 41.46679101859563 | 27.926931604743004 |
| $T_{pre}$ | 0.004932651762315667 | 0.003205401718352722 | 0.0044741068944416425 | 0.0044691432639956475 |
| $T_{post}$ | 0.003421137688008185 | 0.002 | 0.003076731823930066 | 0.002633140569552779 |
| $\tau_{glu}^0$ | 0.0131875344402349 | 0.01 | 0.026167647079173226 | 0.03274728642776609 |
| $\tau_{glu}^{rate}$ | $7.981765401850522 \times 10^{-5}$ | $6.6409090796827 \times 10^{-5}$ | $6.305239365023947 \times 10^{-5}$ | $9.509829357266427 \times 10^{-5}$ |
| $\theta_d$ | 0.002582335829575121 | 0.001 | 0.004140617284273709 | 0.002299701912328601 |

Subsequently, to evaluate the robustness of our results, we calculated the average pattern detection success rate using the optimized parameters on a new set of 100 input spike trains, distinct from the 50 input spike trains used during the BO process. Furthermore, for the 3-bit-HP model, where synaptic interactions via HP highly depend on the spike trains received by synapses within the virtual compartments, we averaged the success rates over 100 random seeds used for generating virtual compartments. The results are listed in Table 8. The success rates dropped 1% to 14% across all models and setups compared to the BO results, suggesting that the parameters obtained via the BO process were overfitted to the specific input spike trains used during the optimization phase. However, the general trend remained consistent (the synaptic efficacy discretization reduced the performance while the HP mitigated this loss).

**Table 8:** Results of the average success rate across different models and setups. Each value represents the success rate calculated using the parameters obtained via Bayesian optimization (BO), derived over 100 trials with input spike trains generated using different random seeds. For the 3-bit-HP synapse model, results were additionally averaged over 100 random seeds for virtual compartments.

|  | 64-bit synapse | 3-bit synapse | 3-bit synapse with HP |
| --- | --- | --- | --- |
| Setup 1 | 88% | 49% | 89% |
| Setup 2 | 82% | 42% | 77% |
| Setup 3 | 84% | 39% | 81% |
| Setup 4 | 49% | 1% | 17% |

### 3.2. Heterosynaptic plasticity accelerates the process by which synapses critical for pattern detection reach their maximum synaptic efficacy

To investigate the effect of the HP model on the learning time, we examined the time evolution of synaptic efficacy in each of the three models. As the index of the learning time, we used the “latency to max efficacy”, the time required for all the “relevant” synapses to reach the maximum efficacy (0.857 for the 64-bit synapse model and 7 for the 3-bit synapse model). Here, “relevant” means that the synapse reached the maximum efficacy in a trial. Thus, the relevant synapses depend not only on the input spike trains but also on the random seed used to generate the virtual compartments. Similar to the average pattern detection success rate in the previous section, we calculated the average latency to max efficacy for successful trials using 100 random seeds for generating the input spike trains in the 64-bit and 3-bit models and 100 random seeds for input spike trains combined with 100 random seeds for virtual compartments in the 3-bit-HP model. Table 9 lists the average latency to max efficacy for each model. The cell for Setup 4 with the 3-bit synapse model is marked as NA because its pattern detection success rate is extremely low (1%). We can observe that the average latency to max efficacy is longer in the 64-bit synapse model than in the 3-bit synapse model across all setups. This is presumably because, compared to the 3-bit synapse model, the 64-bit synapse model has a finer granularity of synaptic efficacy updates per plasticity induction, resulting in smaller changes, thereby requiring more time for the synaptic efficacy to reach the maximum value. By incorporating HP into the 3-bit model, the latency to max efficacy further decreased in all considered setups. This reduction may be due to the suppression of synapses that are irrelevant to the pattern detection in the early stages by HP. This reduces post-synaptic spikes outside of pattern presentation, thereby decreasing the probability that the synapses relevant to the pattern detection undergo LTD. Table 10 lists the coefficient of variation (*CV* = *σ*/*µ*) of the average latency to max efficacy. Here, it is also marked as NA for Setup 4 with the 3-bit synapse model owing to an extremely low success rate (1%). In the case of the 64-bit synapse model, the CV of the average latency to max efficacy remained stable at approximately 16%. By contrast, in the 3-bit synapse and 3-bit synapse with HP models, the CV reached approximately 95%, indicating an extremely large variation relative to the average. This suggests that the average latency to max efficacy becomes highly unstable from trial to trial owing to the reduced bit resolution of synaptic efficacy. This instability is attributed to the following factors associated with low-bit resolution. If correct learning occurs in many synapses immediately after learning begins, the important synapses for pattern detection reach maximum efficacy rapidly. Conversely, if incorrect learning occurs in certain synapses immediately after the start of learning, it requires considerable time to relearn correctly because the change in efficacy induced by a single plasticity event is coarse. Therefore, the value of latency to max efficacy fluctuate greatly depending on each input spike train. The disadvantageous scenario can be addressed by using multiple neurons in applications.

**Table 9:** Average latency to maximum efficacy across different models and setups. The latency to maximum efficacy was defined as the time required for all relevant synapses to reach their maximum efficacy. Each value represents the average over successful trials using 100 random seeds for input spike trains (with an additional 100 seeds for virtual compartments in the 3-bit-HP synapse model). NA indicates that the average could not be reliably calculated because the pattern detection success rate was extremely low (1%).

|  | 64-bit synapse | 3-bit synapse | 3-bit synapse with HP |
| --- | --- | --- | --- |
| Setup 1 | 112.5 s | 68.97 s | 50.00 s |
| Setup 2 | 132.3 s | 58.73 s | 45.83 s |
| Setup 3 | 132.0 s | 76.51 s | 54.93 s |
| Setup 4 | 137.0 s | NA | 60.13 s |

**Table 10:** Coefficient of variation (CV) of the latency to maximum efficacy across different models and setups. The CV was defined as the ratio of the standard deviation to the mean (*CV* = *σ*/*µ*) of the latency to maximum efficacy. NA indicates that the value could not be reliably determined because the pattern detection success rate was extremely low (1%).

|  | 64-bit synapse | 3-bit synapse | 3-bit synapse with HP |
| --- | --- | --- | --- |
| Setup 1 | 15.9% | 72.1% | 87.1% |
| Setup 2 | 9.10% | 92.4% | 94.6% |
| Setup 3 | 8.46% | 52.4% | 79.3% |
| Setup 4 | 7.10% | NA | 76.8% |

### 3.3. Heterosynaptic plasticity promotes bimodality in synaptic efficacy distributions

Postsynaptic firing in response to the target pattern is induced by specific temporary correlated inputs arriving via a small number of synapses with high synaptic efficacy. In contrast, synapses associated with unrelated input spike trains are maintained at weak levels to ensure signal-to-noise separation. Thus, the bimodality of the synaptic efficacy distribution, which captures this distinct split between strong and weak synapses, appears critical for the high selectivity of the learned repetitive spike pattern. The previous study [9] using pattern detection models with 64-bit synaptic efficacy showed that the distribution of synaptic efficacy became bimodal after learning: a small group of synapses reached the maximum efficacy, whereas the majority reached the minimum. Therefore, we investigated the synaptic efficacy distribution after learning in each model. The average synaptic efficacy distribution using 100 different input spike trains for the 64-bit and 3-bit synapse models was calculated. For the 3-bit synapse with HP model, we used 100 random seeds for input spike trains and 100 random seeds for virtual compartments. Figure 2 shows the results, where the horizontal axis represents the synaptic efficacy and the vertical axis represents the number of synapses with that efficacy. For easy comparison, the results for the 64-bit synapse model are shown discretized to 3-bit resolution. Consistent with previous research [9], in the 64-bit synapse model, most synaptic efficacies were concentrated near 0, and the pattern detection was performed by a small group of synapses with maximum efficacy. In contrast, for the 3-bit synapse model, where the success rate significantly decreased, the number of synapses with efficacy 0 and 7 decreased, whereas the number of synapses with efficacy 1 and 2 significantly increased. This trend was consistent across all setups. This indicates that with low-bit resolution synaptic efficacy, supressing the synapses irrelevant to the pattern detection to the minimum value becomes difficult because the large discrete updates characteristic of the 3-bit model amplify the impact of accidental LTP triggered by noise in a large number of such synapses. It induces significant membrane potential fluctuations and postsynaptic firing at undesirable timing, which hinders the selective potentiation of the few synapses critical for the pattern detection. This characteristic is considered a primary factor in the degradation of the success rate. By contrast, by incorporating HP, the distribution of synaptic efficacy after learning in the 3-bit synapse model approached that of the 64-bit model in all setups. This is presumably because HP shortens the latency to max efficacy, enabling synapses important for pattern detection to rapidly reach their maximum value. Notably, this rapid saturation reduces the time window during which the irrelevant synapses can fluctuate at intermediate levels. The rapid formation of strong synapses allows the immediate and effective suppression of other synapses via HP, thereby ensuring a sharp bimodal distribution by driving unimportant synapses to the minimum value.

**Figure 2:**
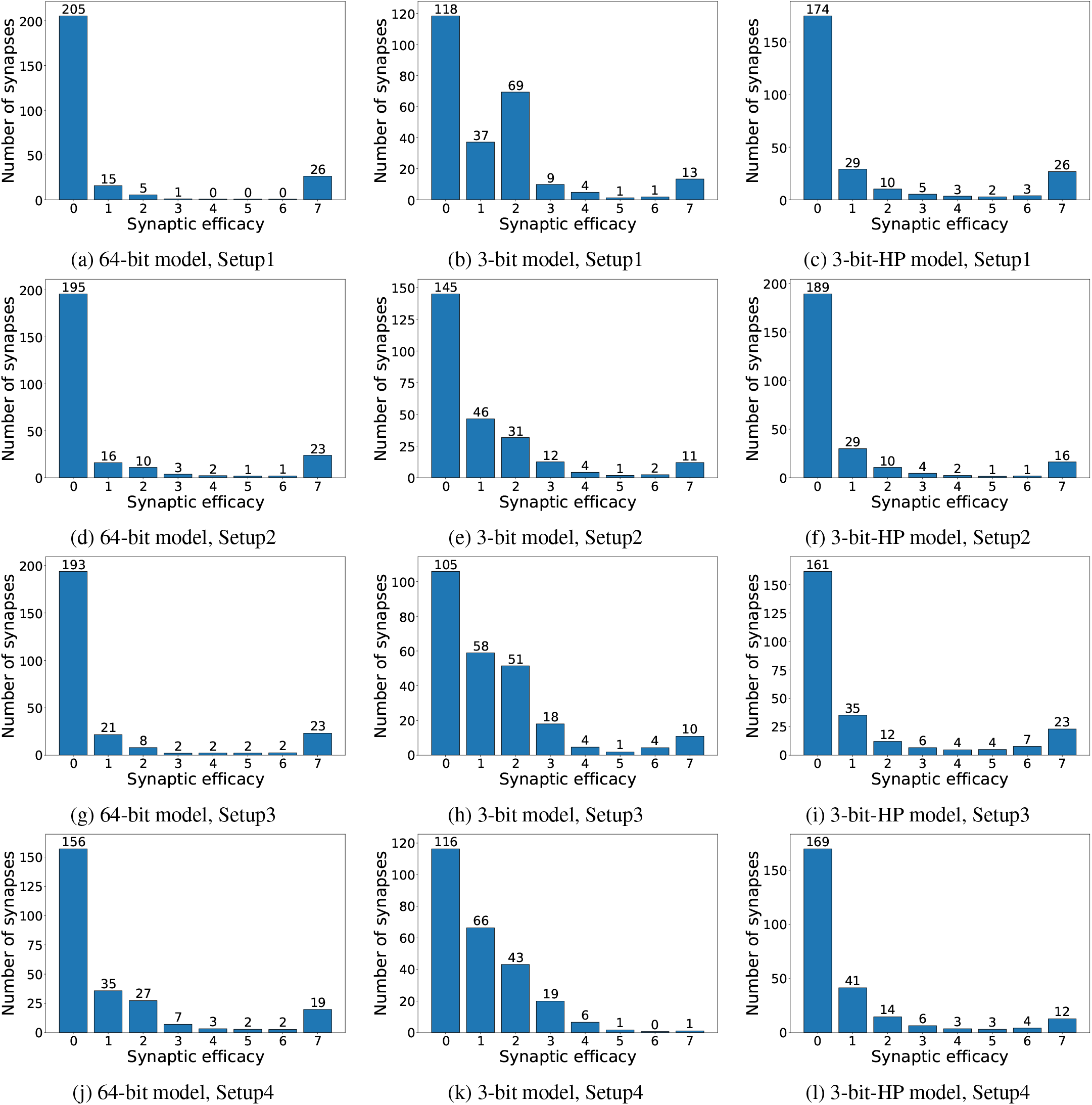
**a–c** Distribution of synaptic efficacy after learning in Setup1. **d–f** Distribution of synaptic efficacy after learning in Setup2. **g–i** Distribution of synaptic efficacy after learning in Setup3. **j–l** Distribution of synaptic efficacy after learning in Setup4.

## 4. Discussion

### 4.1. Impact of the proposed heterosynaptic plasticity on spatio-temporal spike pattern detection with 3-bit synaptic efficacy

In this study, we proposed HP model based on glutamate spillover, whose effect on a spatio-temporal spike pattern detection model with a limited resolution of synaptic efficacy was evaluated. Our results demonstrated that the proposed HP effectively mitigated the performance degradation in spike pattern detection induced by low-bit discretization of synaptic efficacy. Notably, in Setups 1 to 3 (see Tables 4–8), the 3-bit-HP synapse model achieved a pattern detection success rate comparable to that of the 64-bit synapse model. However, in Setup 4, the success rate of the 3-bit-HP synapse model remained significantly lower than that of the 64-bit synapse model. To investigate whether this performance degradation stems from the stochastic nature of the 10 Hz background noise or from the jitter noise, we evaluated the spike pattern detection performance in each model under following two additional setups. The first one is Setup 4a in which the discrete 10 Hz noise spikes in Setup 4 were replaced by an equivalent background current, *V*_bg_(*t*). The second is Setup 4b, which is identical to Setup 4, but with the jitter noise eliminated. The equivalent background current in Setup 4a, *V*_bg_(*t*), was derived as follows. As defined in the Methods section, the EPSP kernel *ϵ*(*s*), which represents the contribution of a single input spike to the membrane potential in the SRM, is as follows:

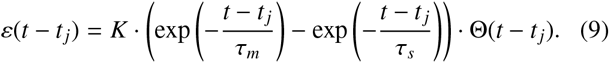

The time integral of this kernel, *A*_*ϵ*_, was calculated as follows.

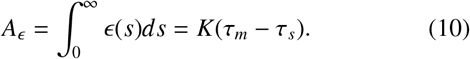

The expected contribution of the *V*_bg_(*t*) is proportional to the sum of all synaptic efficacies at time *t* and is expressed as follows:

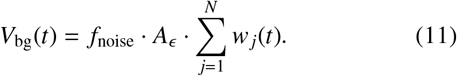

Here, *N* = 256 is the total number of synapses, *w*_*j*_ ∈ [0, 1] represents efficacy of synapse *j*, and *f*_noise_ = 10 Hz represents mean firing rate of the noise. The membrane potential *p*(*t*) in the first setup is expressed by the sum of the *η* kernel, the EPSPs provided by presynaptic spikes, and the background term *V*_bg_(*t*).

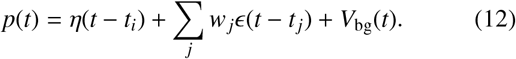

Simulation results are summarized in Table 11. Simulation results revealed that in the 64-bit synapse model, the pattern detection success rates for Setups 4a and 4b were 57% and 58%, respectively, representing an increase of 8% and 9% compared to Setup 4. In contrast, the 3-bit synapse model yielded success rates of 4% for 4a and 14% for 4b, reflecting the improvements of 3% and 13% over Setup 4, respectively. The 3-bit synapse model with HP achieved success rates of 30% (4a) and 27% (4b), marking increases of 13% and 10% relative to Setup 4, respectively. Regarding noise sensitivity, the 64-bit and 3-bit-HP synapse models exhibited comparable degradation under 10 Hz and jitter noise. However, in the 3-bit synapse model, jitter noise was found to be a more significant factor in reducing the success rate than 10 Hz noise. In the 64-bit model, minor fluctuations in spike timing owing to the jitter noise do not substantially alter the magnitude of synaptic efficacy changes induced by STDP. Conversely, the 3-bit model utilizes rectangular STDP, where no plasticity is induced if the spike timing falls outside the defined plasticity window. Consequently, jitter-induced timing shifts are more likely to prevent plasticity induction, increasing the model’s susceptibility to the jitter noise. In the 3-bit-HP synapse model, even if the jitter noise prevents the induction of depression via rectangular STDP, HP can induce LTD. This compensatory mechanism appears to mitigate the impact of the jitter noise. Further investigation is required to fully elucidate the underlying mechanisms governing the observed fluctuations in success rates by noise.

**Table 11:** Results of the average success rate in setups 4a and 4b across different models. Setup 4a represents a condition where 10 Hz noise were replaced by an equivalent background current, *V*_bg_(*t*). Setup 4b is identical to setup 4 but with the jitter noise eliminated. The success rate calculated using the parameters obtained via Bayesian optimization (BO) in setup 4, dirived over 100 trials with input spike trains generated using different random seeds. For the 3-bit-HP synapse model, results were additionally averaged over 100 random seeds for virtual compartments.

|  | 64-bit synapse | 3-bit synapse | 3-bit synapse with HP |
| --- | --- | --- | --- |
| Setup 4a | 57 % | 4 % | 30 % |
| Setup 4b | 58 % | 14 % | 27 % |

Our study demonstrated that the proposed HP reduced the latency to max efficacy. Given that this HP mechanism exclusively induces LTD in our model, this result appear counterintuitive. However, this phenomenon can be explained by the depression of synapses irrelevant to pattern detection during the early stages of learning. By suppressing these irrelevant synapses, HP facilitates the occurrence of postsynaptic spikes at a timing in the target pattern presentations more effectively than in models without this mechanism. Consequently, this reduces the likelihood of inducing unintended LTD via STDP at relevant synapses for pattern detection. Quantitative verification of this hypothesis remains a subject for future investigation.

Furthermore, analysis of the synaptic efficacy distribution after learning revealed that HP reduced the number of synapses with intermediate efficacies while increasing the count of synapses reaching maximum efficacy. This suggests that the synapses relevant to the pattern detection actively suppress their neighboring irrelevant synapses, thereby promptly correcting unfavorable LTPs. Our HP model suppresses the synapses irrelevant to pattern detection, thereby reducing postsynaptic firing outside the target pattern. This mechanism enhances the likelihood of postsynaptic spikes during the target pattern. Thus, the risk of accidental LTD via STDP at relevant synapses is mitigated, enabling them to achieve maximum synaptic efficiency. Collectively, all of the aforementioned results suggest that the proposed mechanism promotes a bimodal distribution of synaptic efficacy, which enhances selectivity for the target pattern, sustaining high pattern detection performance despite low-resolution synaptic efficacy.

### 4.2. Physiological basis of the proposed model

The primary motivation of this study is to build an advanced neuromorphic learning model by drawing inspiration from neurophysiological findings that have been largely overlooked in existing neuromorphic models. The second motivation is to contribute to elucidating the brain microciruit’s information processing by the “analysis by construction” approach. Consequently, considering the biological plausibility of the proposed model is crucial. Our model assumed that glutamate spillover activates NMDARs on the adjacent spines. When this activation is combined with bAP-induced potential elevation, it may trigger calcium influx and subsequent heterosynaptic LTD. The number of synapses affected by glutamate spillover, defined here as the number of synapses within a virtual compartment, remains experimentally unclear. This value is believed to depend primarily on the spine density and the extent of glutamate diffusion. In this study, we assumed that each virtual compartment contains 6 to 10 synapses. According to a previous study [23], spine densities vary across brain regions: 1.14 ± 0.723 spines/*µ*m in the somatosensory cortex and 3.04 ± 0.825 spines/*µ*m in the hippocampal CA1 proximal stratum radiatum (PSR) [23]. In the CA1 region, density also depends on the dendritic location, with the stratum lacunosum-moleculare (SLM) exhibiting a lower density of 0.75 ± 0.360 spines/*µ*m than the PSR. Although the precise diffusion distance of glutamate in vivo remains unclear, studies using acute hippocampal CA1 slices have shown that glutamate from a single quantal release can reach distances exceeding 1.5 *µ*m and activate receptors [24]. Diffusion is further enhanced during multiquantal release triggered by a single action potential. Specifically, the release of five vesicles can activate NMDARs approximately 2.0 *µ*m away. This implies that spillover can influence 8–16 synapses in the CA1 PSR and 2–4 synapses in the SLM. The number of synapses per virtual compartment assumed in this study was selected assuming similar orders of magnitude in the other cortex region. Furthermore, in the somatosensory cortex, extrasynaptic glutamate from spillover is known to persist at micromolar concentrations for tens of milliseconds [13]. This concentration is sufficient to activate NMDARs but insufficient for *α*-amino-3-hydroxy-5-methyl-4-isoxazolepropionic acid receptors (AMPARs), justifying our model’s assumption that glutamate spillover exclusively affects NMDARs.

### 4.3. Future works

In this study, we used an I&F-based point neuron model, which ignores two critical features of the neuronal cells: complex firing patterns such as bursting spike at soma, and spike events on the dendrites (dendritic spikes). Ignoring these factors limits the scope of our proposed HP on the distal dendrites because backpropagating action potentials (bAP) amplitude decreases as it propagates away from the soma. To consider the plasticity at distal dendrites bAP has to be modeled because it is required to remove the Mg^2+^ block of NMDARs during periods of high glutamate concentration in extrasynaptic space. This bAP attenuation might be compensated by somatic burst firing or dendritic spikes, which provide the necessary depolarization to the distal dendrites. Incorporating these factors into the model is a crucial future direction.

The synaptic efficacy depends on both the amount of presynaptic glutamate release and the number of postsynaptic AMPARs. Previous research [25] indicates that glutamate spillover can induce heterosynaptic depression by activating presynaptic metabotropic glutamate receptors (mGluRs), which reduces subsequent glutamate release. This mechanism could be beneficial when a virtual compartment contains numerous relevant synapses for pattern detection that should be protected from the depression process in our HP model. By reducing the amount of spillover via presynaptic inhibition[25], the induction of our HP could be suppressed. Introducing heterosynaptic LTP may be pivotal for improving the success rate of pattern detection. Because if a virtual compartment contains many relevant synapses for pattern detection, heterosynaptic LTP can collectively potentiate them. However, because our model treats pre- and post-synaptic efficacies with a single parameter, it may lead to a positive feedback loop: increased synaptic efficacy leads to greater glutamate spillover per spike, which facilitates further heterosynaptic LTP. The aforementioned presynaptic mGluR-mediated inhibition of glutamate release could serve as a potential solution to stabilize this feedback. These remain subjects for future investigation.

We demonstrated that the performance of a spatiotemporal spike pattern detection task with low-resolution synapses can be boosted by a learning model inspired by HP. For hardware implementation, this model requires a virtual compartment circuit for 6 to 10 synapses. Thus, efficient implementation has to be pursued not only by developing optimized circuit architecture but also by adapting the model to the characteristics of the electronic devices. It is the most important future work.

